# PIEZOs regulate oligodendrocyte sheath formation, expansion, and myelination potential

**DOI:** 10.64898/2026.04.23.720488

**Authors:** AM Coombs, D Heo, DJ Orlin, CL Call, ME Bechler, SE Murthy, B Emery, KR Monk

**Affiliations:** Vollum Institute, Oregon Health & Science University, Portland, OR, USA; Jungers Center for Neurosciences Research, Department of Neurology, Oregon Health & Science University, Portland, OR, USA; Department of Cell and Developmental Biology, State University of New York Upstate Medical University, Syracuse, NY, USA; Department of Neuroscience and Physiology, State University of New York Upstate Medical University, Syracuse, NY, USA

**Author notes:** Co-corresponding authors: Dr. Ben Emery and Dr. Kelly R. Monk. Equal contributing authors (D Heo, DJ Orlin). National Center for Complementary and Integrative Health, National Institutes of Health (DJ Orlin).

## Abstract

Myelination requires precise integration of physical cues by oligodendrocyte lineage cells (OLCs), but the molecular sensors that detect these cues remain incompletely understood. Here, we demonstrate that oligodendrocyte progenitor cells (OPCs) are sensitive to sub-micron changes in membrane displacement. Based on channel properties, RNA expression, and protein abundance, we find that the mechanosensitive ion channel PIEZO1 contributes to OPC mechanosensitivity. *In vivo*, zebrafish with oligodendrocyte (OL)-specific disruption of *piezo1* have fewer sheaths per OL. Zebrafish with OL-specific *piezo2* disruption also have fewer sheaths as well as decreased total myelin capacity over time. OL-specific disruption of both *piezo1* and *piezo2* caused more severe phenotypes, with reduced OPC volume, and in myelinating OLs, reduced sheath number, sheath length, and total myelin output. Furthermore, *piezo1*/*piezo2* disruption leads to sporadic sheath formation outside the normal developmental window. Our findings indicate that OLs use Piezo channels *in vivo* to influence sheath formation, expansion, and retractions.

## INTRODUCTION

The rapid transmission of neuronal signals in jawed vertebrates requires the ensheathment of axons by myelin, which is generated by oligodendrocytes (OLs) in the central nervous system (CNS). Myelination begins when morphologically complex and highly dynamic OL precursor cells (OPCs) differentiate, first into premyelinating OLs and finally into mature OLs capable of myelinating multiple axon segments. During this process, OLs undergo profound morphological changes as their cellular extensions ensheathe axons and begin to form sheets of myelin. The leading edge of the OL process circumnavigates the axon and pushes underneath previous wraps, allowing for expansion of the membrane both longitudinally and in thickness^1^. Mechanical force is thought to be required to displace myelin from the axon to make way for the formation of new layers of membrane, which is generated by actin filament turnover at the leading edge^2,3^. Contact and mechanical interactions between oligodendrocyte lineage cells (OLCs) and axons are persistent throughout the myelination process and may serve as a key point of bidirectional information exchange between neurons and OLs. The molecular sensors and mechanisms that OLs use to gather information from the environment to inform features of myelination, such as the number of sheaths, sheath length, and the overall myelinogenic potential of a cell, remain underexplored.

Myelin output can vary by OL, characterized by differences in sheath number, length, and total myelination potential with regional heterogeneity in the CNS^4^. Irrespective of these morphological differences, OLs maintain similar conductance when normalized for the number and length of processes^5^. This suggests that the morphological variability in OL populations may simply reflect their ability to sense and respond to environmental differences between regions in the CNS, such as axon size, density, and the demands of the local circuitry. OLs have been shown to sense physical properties of axons and scale myelin output accordingly based on their ability to modulate myelin coverage in response to changes in diameter on inert nanofibers that structurally mimic axons^6^ and manipulations that alter the availability of axons *in vivo*^7^. Mechanosensitive ion channels (MSICs) are ideal candidates to serve as sensors of the extracellular environment because they are specialized proteins that respond to mechanical cues and convert them into biological signals in the form of the flow of ions across a membrane^8^. Cationic MSICs are of particular interest because of the well-established role for calcium signaling in regard to the regulation of sheath extensions and retractions^9–11^.

MSICs are membrane proteins that possess multiple transmembrane helices and a pore-forming domain gated by the application of physical force. MSICs are widely expressed in a variety of cell types, and there is a growing appreciation for their role in the regulation of a range of biological processes, including in the nervous system. For example, MSICs in neurons that innervate the skin mediate the sense of touch in mice and humans^12–14^, and in microglia allow for the detection and clearance of Aβ plaques^15^. Of particular interest, rodent OLCs have been reported to express genes encoding members of the PIEZO and TMEM63 families of MSICs, including *Piezo1*, *Piezo2*, and *Tmem63a*^16–18^. In one dataset, expression of *Tmem63a* is highest in newly formed and mature OLs, while *Piezo1* is highest in OPCs with decreased abundance as OLs differentiate^16^. Several recent publications investigate the role of TMEM63A in CNS myelination^19–22;^ however, the relative roles of PIEZOs in OLs remain underexplored. PIEZOs were the first identified family of ion channels required for mammalian mechanosensation^8,23^. Whereas PIEZO1 and PIEZO2 typically have nonoverlapping expression and function, a small number of cell types (*e.g.*, subpopulation in somatosensory ganglia and chondrocytes) co-express both channels^24^. Notably, OLCs have been reported to express both *Piezo1* and *Piezo2*^17^, creating an opportunity for these channels to have independent functions, compensate for one another, or work in synergy to regulate the OL lineage. The independent and combined functions of PIEZOs in the OL lineage remain incompletely understood. It was previously reported that PIEZO1 mediates an OPC’s ability to respond to substrate stiffness under pathogenic conditions and in normal aging^25^. More recently, roles for PIEZO1 in activity-dependent OPC proliferation and glioma progression as well as axon diameter sensing by OLs have been described^26,27^, altogether supporting the notion that OLCs use mechanical cues from the environment to shape multiple aspects of their biology.

Here, we use electrophysiology to characterize the mechanosensitivity of OLCs, which revealed that OPCs have the capacity to respond to sub-micron changes in membrane displacement. We explore the role of PIEZO1 in the cell-autonomous regulation of the OL lineage using primary rodent cultures plated onto inert nanofibers, which reveal reduced sheath length and total myelin output per cell, suggesting that the ability to sense mechanical cues via PIEZO1 influences myelination. Lastly, we investigate the independent and combined contributions of Piezo1 and Piezo2 *in vivo* during developmental myelination using cell-type-specific knockdown approaches in zebrafish. These experiments uncovered that PIEZOs have independent functions within the OL lineage, and double knockdown revealed altered sheath number, length, myelination capacity, and sheath dynamics. In addition to contributing to the understanding of the basic biology of how OLs sense and respond to their environment, the identification of new mechanisms of regulating OL myelination could reveal novel pathways and therapeutic targets for the treatment of demyelinating diseases, leukodystrophies, and other white matter deficits of the CNS.

## RESULTS

### OPCs have mechanically-activated (MA) currents in response to membrane displacement

To determine whether OLCs are mechanosensitive, mechanically-activated (MA) currents were recorded from cultured primary rat OPCs immunopanned between postnatal day (P)6-8 using the whole-cell patch clamp configuration. MA currents were induced by 0.4 µm step-wise increases in membrane displacement with a blunt glass probe with 300 ms-long stimulation sweeps (**Fig. 1A**). OPCs showed an average Imax of 503.8 ± 183 pA (**Fig. 1B**; mean ± SE). 13 of the 14 cells tested had a robust MA response with inactivation kinetics of 37.3 ± 2.75 ms (**Fig. 1C**; mean ± SE). These observations demonstrate that OPCs have large, rapidly inactivating currents in response to whole-cell indentation stimulations that strongly resemble PIEZO-mediated currents (*i.e.*, large, rapidly inactivating currents that scale with changes in force on the membrane^8^). Previous studies have shown that indentation-evoked PIEZO1 and PIEZO2-mediated currents are very similar in threshold, rise-time, and channel conductance^8,28,23^, making it challenging to use channel properties alone to identify which channel mediates the mechanosensitivity of OPCs.

**Figure 1:**
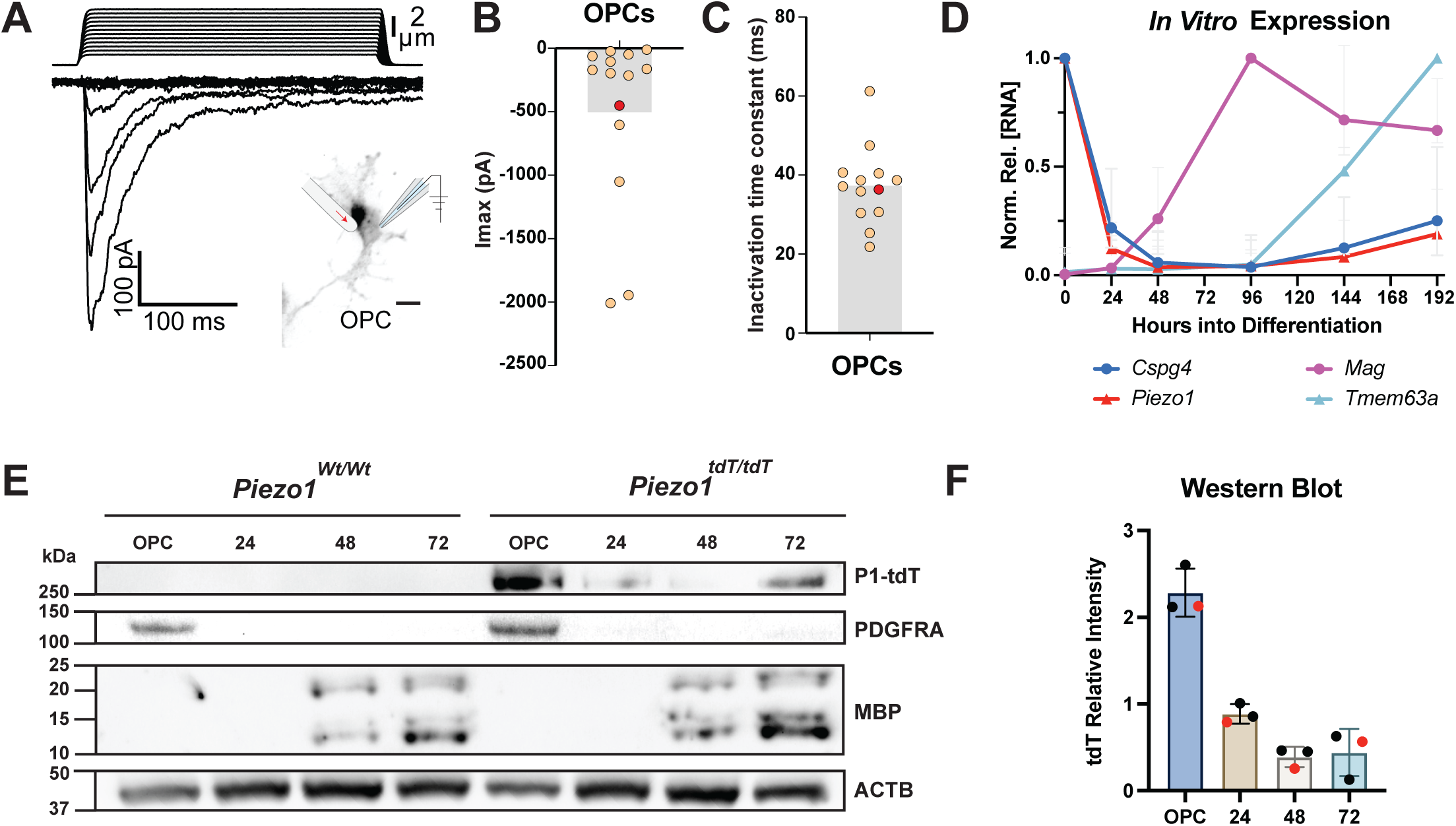
OPCs possess MA currents and express PIEZO1. A) Representative trace from whole-cell patch clamp electrophysiology of a primary rat OPC in culture. The top of the graph is the stimulation trace showing the 300 ms stimulation trace with 0.4 µm step-wise increases in membrane displacement using a glass stimulation rod. The bottom portion of the graph shows the current (pA) trace at each stimulation. 13 of the 14 tested cells showed detectable MA currents. Scale bar: 10 μm B) Maximal current (Imax) for each cell in response to indentation stimulation N = 2 animals, 14 cells, 2 independent cell isolations. The red data point is associated with the representative trace in Fig. 1A. C) Inactivation time constant for each cell responsive to indentation stimulation N = 2 animals, 13 cells, 2 independent cell isolations. The red data point is associated with the representative trace in Fig. 1A. D) qRT-PCR of *Piezo1*, *Tmem63a*, *Mag*, and *Cspg4* in cultured rat OLs during differentiation (0-192 h). Values are normalized to the transcript’s highest expression value during differentiation N = 3 animals, 3 independent cell isolations. E) Western blot analysis of PIEZO1-tdTOMATO, PDGFRA, MBP, and ACTB expression in *Piezo1^tdT/tdT^* and *Piezo1^Wt/Wt^* control murine OPCs and OLs at 24-72h differentiation. F) Quantification of tdTOMATO intensity relative to ACTB in western blots. N = 3 animals per genotype, 3 independent cell isolations. Red dots indicate the data points corresponding to the example image in E.

### OPCs express the mechanosensitive ion channel PIEZO1

To further investigate expression of candidate mechanosensors across OL differentiation, OPCs were isolated from P6-8 rats by immunopanning and either maintained in a proliferative state in the presence of platelet-derived growth factor (PDGF) or differentiated into OLs by the removal of PDGF and replacement with thyroid hormone (T3). RNA was isolated from cultured OPCs and at various time points into differentiation, ranging from 24 to 192 hours. Given that the OPC MA currents resemble PIEZO-like currents and that TMEM63A is an important regulator of myelination^19–22^, we focused on examining *Piezo1*, *Piezo2*, and *Tmem63a* expression profiles in OPCs, using *Cspg4* as a marker for the OPC stage and *Mag* for postmitotic OLs. As anticipated, *Cspg4* was most highly expressed in the OPC population, and *Mag* increased in expression until a plateau around 96 hours. The expression of *Piezo1* closely resembled that of *Cspg4,* with highest expression in OPCs and decreased expression with differentiation (**Fig. 1D**). In contrast, *Tmem63a* did not show detectable expression until 144 hours into differentiation (**Fig. 1D**). *Piezo2* was not detected in OLC cultures through the first 196 hours into differentiation; however, qPCR probes were able to verify expression in DRGs. The expression profile of these candidate MSICs, in combination with the large currents and rapid inactivation kinetics seen in the electrophysiology experiments performed on OPCs, led us to focus on Piezo1.

Due to a lack of reliable antibodies, a direct assessment of PIEZO1 localization and protein abundance is challenging. To circumvent this, OPCs were isolated from the *Piezo1*-tdTomato mouse line, which features a knock-in of tdTomato into the endogenous *Piezo1* allele, leading to a fusion of tdTOMATO to the C-terminus of PIEZO1^29^. Wild-type or *Piezo1-tdTomato* (*Piezo1^tdT^*) OPCs were plated into proliferation media or differentiation media. Protein was harvested from OPCs at 24, 48, and 72 hours into differentiation and western blot was performed using anti-RFP, anti-PDGFRA, anti-MBP, and anti-ACTB antibodies (**Fig. 1E**). Consistent with the above RT-qPCR results, western blots revealed that PIEZO1 protein abundance is highest in PDGFRA+ OPCs, with a decreased abundance as differentiation proceeds and MBP levels increase (**Fig. 1F**). To determine whether OLCs showed heterogenous levels of PIEZO1 during differentiation, immunopanned *Piezo1^tdT^* and wild-type cells were plated onto coverslips and maintained as OPCs in proliferation media or were plated into differentiation media for 48 hours. Cells were stained with anti-RFP to detect PIEZO1, anti-PDGFRA, and anti-MBP (**Supplement Fig. 1A-B**). The majority of PDGFRA+ OPCs (88.6%) showed detectable tdTomato signal, whereas it was detectable in only 7.2% of MBP+ OLs at 48 hours into differentiation (**Supplement Fig. 1C**). These expression results, coupled with the electrophysiology experiments, suggest PIEZO1 as a likely candidate to mediate the mechanosensitivity of OPCs.

### PIEZO1 is not required for OPC proliferation or differentiation *in vitro*

To investigate PIEZO1’s contributions to the cell-autonomous regulation of OL lineage progression and myelination, *Piezo1* was genetically removed from OLCs in late embryonic development. *Piezo1^Floxed^* mice^30^, in which exons 20-23 of the *Piezo1* gene are loxP flanked, were crossed with an *Olig2^Cre^* line^31^ to generate *Piezo1^CKO^* mice (*Piezo1^Fl/Fl^;Olig2^Cre/+^*). To account for any potential effects of *Olig2* haploinsufficiency, *Piezo1^Wt/Wt^;Olig2^Cre/+^*littermate animals were used as controls. To verify removal of *Piezo1* from the OL lineage, calcium imaging was performed using the ratiometric calcium indicator Fura2 in the presence of the highly-selective PIEZO1 agonist, Yoda1^32^. Control cells showed a striking rise in intracellular calcium upon the application of Yoda1, further confirming high Piezo1 expression in OPCs, whereas *Piezo1^CKO^* cells do not respond to pharmacological activation of PIEZO1 (**Fig. 2A-C, Video 1-2**). *Piezo1^CKO^* cells showed a 94% reduction in the area under the curve after the application of Yoda1 in *Piezo1^CKO^* cells versus wild-type controls (**Fig. 2D**; Control – N=3 animals, 971 cells, 3 independent cell isolations; CKO – N=3 animals, 1,043 cells, 3 independent cell isolations). In contrast, both genotypes showed a robust rise in intracellular calcium in response to potassium chloride (KCl)-mediated depolarization (**Fig. 2A-C**). The lack of response in *Piezo1^CKO^* cells demonstrates the effective removal of *Piezo1* from the OL lineage.

**Figure 2:**
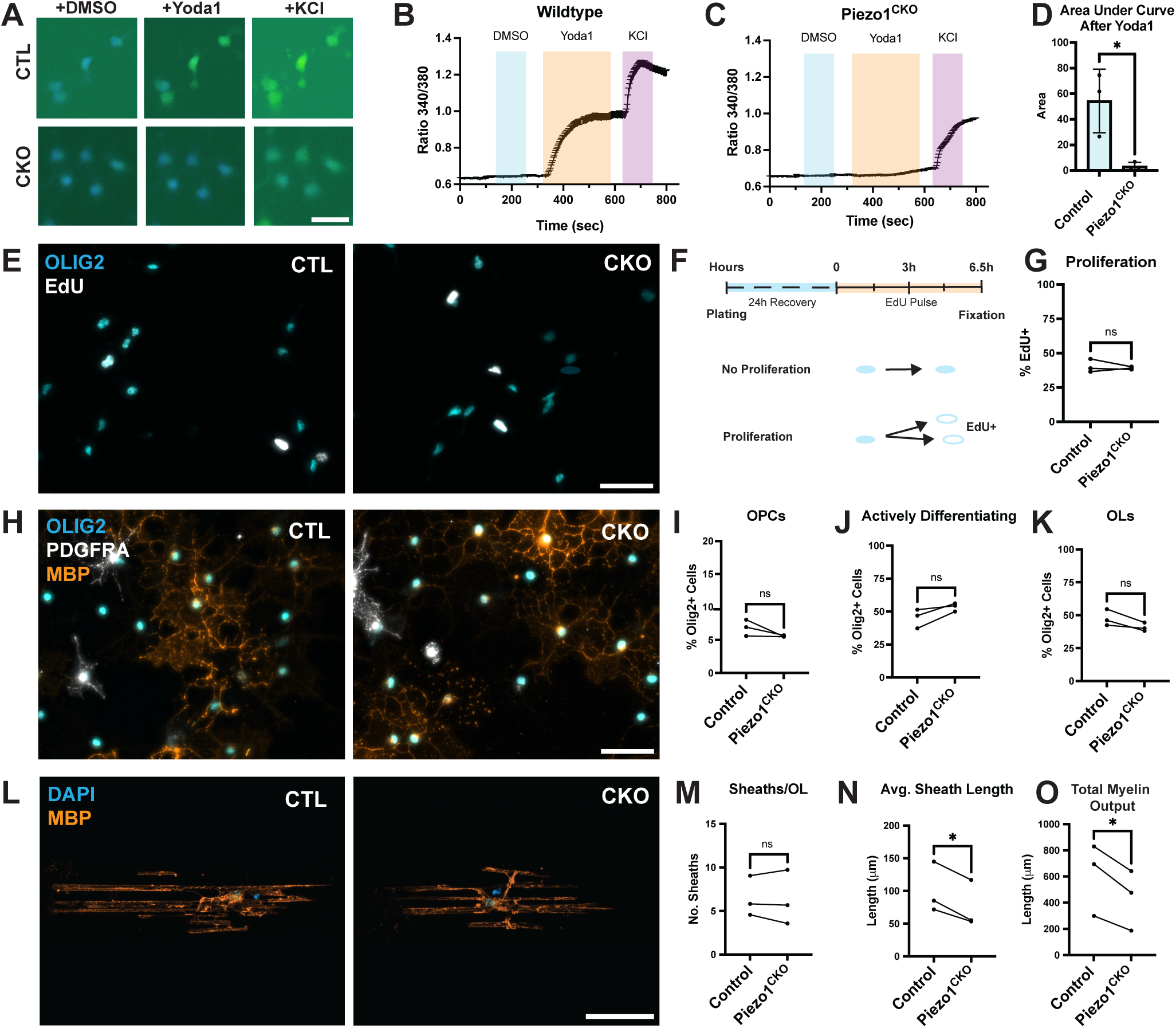
Loss of PIEZO1 does not alter OPC proliferation or differentiation but reduces sheath expansion and total myelin capacity *in vitro*. A. Raw images from Fura2 calcium imaging after the application of DMSO, 30 µM Yoda1, and 200 mM KCl. Scale bar: 25 μm B. Representative calcium imaging trace from control (*Piezo1^Wt/Wt^;Olig2^Cre/Wt^*) OPCs loaded with Fura2 and sequentially exposed to DMSO, 30 µM Yoda1, and 200 mM KCl. Trace averaged from 72 cells imaged from 1 coverslip. C. Representative calcium imaging trace from *Piezo1^CKO^* cells (*Piezo1^Fl/Fl^;Olig2^Cre/Wt^*) loaded with Fura2 and exposed to DMSO, 30 µM Yoda1, and 200 mM KCl. Trace averaged from 121 CKO cells from a single coverslip. D. Quantification of area under the 340/380 curve for each animal during application of Yoda1. The imaging data from 3 coverslips per animal were pooled for analysis for 3 animals per genotype. Data shown as mean ± SEM. p = 0.024 by student’s t-test, N = 3 animals per genotype, 971 control cells and 1043 cells from *Piezo1^CKO^* cells, 3 independent isolations. E. Representative images from EdU proliferation assay of control or *Piezo1^CKO^* cells. OLIG2 is a marker of OLCs, and EdU labels cells that divided over the 6.5-hour time course. F. Experimental design for the detection of proliferating cells *in vitro*. G. Quantification of the proportion of OLIG2+ cells labelled for EdU. N = 3 independent OPC isolations per genotype, with 818 cells assessed for controls and 947 cells for *Piezo1^CKOs^.* p = 0.643 by paired t-test. H. Representative images from control or *Piezo1^CKO^* cells fixed 48 hours into differentiation and stained for OLIG2, PDGFRA, and MBP. I. Quantification of the percentage of OLIG2+ cells that are PDGFRA+ OPCs. Control experimental totals: 859 cells, 3 animals, 3 independent OPC isolations CKO experimental totals: 824 cells, 3 animals, 3 independent OPC isolations. p = 0.207 by paired, 2-tailed t-test. J. Quantification of the percentage of OLIG2+ cells that are actively differentiating (PDGFRA-, MBP-). Control experimental totals: 859 cells, 3 animals, 3 independent OPC isolations CKO experimental totals: 824 cells, 3 animals, 3 independent OPC isolations. p = 0.101 by paired, 2-tailed t-test. K. Quantification of the percentage of OLIG2+ cells that are MBP+ OLs. Control experimental totals: 859 cells, 3 animals, 3 independent OPC isolations CKO experimental totals: 824 cells, 3 animals, 3 independent OPC isolations. p = 0.097 by paired, 2-tailed t-test. L. Representative images of a control and Piezo1^CKO^ cell plated onto nanofibers and allowed to differentiate for 7 days before being fixed and stained with DAPI and MBP. M. The number of sheaths per OL for Control and *Piezo1^CKO^* cells. p = 0.767 by paired, 2-tailed, t-test. N. Average sheath length per OL for Control and *Piezo1^CKO^* cells. p = 0.019 by paired, 2-tailed t-test. O. Average myelin output (summed sheath lengths per OL) for control (*Piezo1^wt/wt^;Olig2^Cre/wt^*) and *Piezo1^CKO^* cells. p = 0.032 by paired, 2-tailed t-test. For M-O, Control experimental totals: 50 cells, 3 animals, 3 independent cell isolations. CKO experimental totals: 48 cells, 3 animals, 3 independent cell isolations. For E, H, and L – Scale bar: 50 μm

To determine whether loss of *Piezo1* altered OPC proliferation, OPCs were isolated from *Piezo1^CKO^* and littermate controls, plated onto coverslips, and given 24 hours to recover before a 6.5-hour pulse with EdU (5’-ethynyl-2’-deoxyuridine) (**Fig. 2E-F**). The proportion of dividing cells [(OLIG2+ EdU+)/OLIG2+] was quantified across three independent cell isolations with three animals per genotype (**Fig. 2G**). There was no significant difference in proliferation rates between control and *Piezo1^CKO^* cells. To determine whether loss of *Piezo1* alters OPC differentiation, cells were switched from proliferation to differentiation media and allowed to incubate for 48 hours before fixation. Cells were next stained with anti-OLIG2, anti-MBP, and anti-PDGFRA (**Fig. 2H**), classified, and quantified based on their immuno-positivity for these markers of various stages of OL development. There was no significant difference in the ratio of OLIG2+, PDGFRA+ OPCs (**Fig. 2I**), actively differentiating OLIG2+, PDGFRA-, MBP- (**Fig. 2J**), or mature OLIG2+, MBP+ OLs (**Fig. 2K**) between *Piezo1^CKO^* and control cells. In all, our data suggest that in the context of these two-dimensional cultures, the cell-autonomous regulation of OPC proliferation and differentiation is unaltered by the genetic deletion of *Piezo1*.

### *Piezo1^CKO^* cells have reduced sheath expansion and total myelin capacity *in vitro*

While 2D culture enables the detection of proteins expressed by mature OLs, it fails to evaluate key morphological features of myelination, such as sheath number, average sheath length, or total myelin output per cell. To determine whether PIEZO1 contributes to the cell-autonomous regulation of sheath formation, OLCs were plated onto 3D polystyrene nanofibers, which mimic the mechanical features of axons without the molecular signals contributed by neurons^33^. Mouse OPCs were immunopanned between P6-8 and were expanded before being plated onto 3D nanofiber scaffolds and allowed to differentiate for 7 days. Cells were then stained using DAPI and anti-MBP to label OL sheaths before being mounted onto glass slides as previously described^34^. Cells were reconstructed (**Fig. 2L**) to assess the number of sheaths per OL, the average sheath length, and the total myelin output per cell. Total myelin output is the sum of the sheath length per OL and is a metric of the myelination capacity of each cell^35^. There was no difference in the number of sheaths formed per OL between control and *Piezo1^CKO^* cells (**Fig. 2M**); however, *Piezo1^CKO^* cells showed a significant reduction in average sheath length (**Fig. 2N**) and total myelin output (**Fig. 2O**) per cell compared to controls.

These results indicate that the loss of *Piezo1* does not alter proliferation, differentiation, or sheath formation. However, knockout of *Piezo1* from the OL lineage leads to a decrease in average sheath length and total ensheathment capacity per cell when plated onto nanofibers. This suggests that the ability of OLCs to sense mechanical cues through PIEZO1 results in a cell-autonomous regulation of sheath expansion and the total myelin capacity of a cell, raising the question whether a similar regulatory process is occurring *in vivo*.

### Knockdown of *piezo1* in OLCs reduces myelin formation *in vivo*

In order to visualize OLCs with single-cell resolution and track them longitudinally throughout development *in vivo*, we turned to cell-type-specific experiments in zebrafish. *Piezo1* is expressed in a variety of cell types and has been shown to play an important role in axon targeting and vascular development^29,36^, precluding the use of global knockouts for the assessment of a cell-autonomous role in myelination. We therefore targeted *piezo1* mRNA using a plasmid-based CRISPR-Cas13d system; when paired with the UAS/KalTA4 binary expression system, this allows for cell-type-specific gene knockdown (KD) in zebrafish OLs (Heo et al., submitted). By injecting the KD plasmid into *tg(uas:mScarlet;sox10:kalta4)* zebrafish, targeted cells (eGFP+) and non-targeted control cells (mScarlet+) can be visualized within the same animal and tracked from 3 to 7 days post-fertilization (dpf), spanning zebrafish developmental myelination. Based upon our *in vitro* findings, we hypothesized that changes in OL sheath expansion and total myelin capacity would be observed.

Two crRNAs targeting *piezo1* were cloned into the targeting vector *ptol2-UAS-RfxCas13d-P2A-EGFP-CAAX;U6x-piezo1* (**Fig. 3A**). Embryos were collected from *tg(uas:mscarlet;sox10:kalta)* fish and injected with the all-in-one plasmid. Zebrafish were longitudinally imaged at 3, 5, and 7 dpf (**Fig. 3B**), which allowed cells to be grouped based on the time of initial sheath formation and provided a means to compare features of interest across time in cells of a similar age^37^ (**Supplement Fig. 2**). The day of initial sheath formation was referred to as OL1. Therefore, cells with sheath formation on the first day of imaging were tracked through OL5. Cells that were identified as OPCs were tracked from OL1-3. Myelin sheaths were traced using Simple Neurite Tracer in Fiji to quantify the number of sheaths, average sheath length, and total myelin output per OL. Although non-targeted controls and *piezo1-KD* cells showed similar developmental trajectories (**Fig. 3C**), a significant reduction in the number of sheaths per OL was observed in *piezo1-KD* cells relative to non-targeted controls across all stages of development examined (OL1-5, **Fig. 3D**). However, when comparing average sheath length and total myelin output, no difference was observed (**Fig. 3E-F**). These data suggest that Piezo1 regulates myelin sheath formation *in vivo* but does not significantly impact sheath expansion or total myelin capacity of a cell.

**Figure 3:**
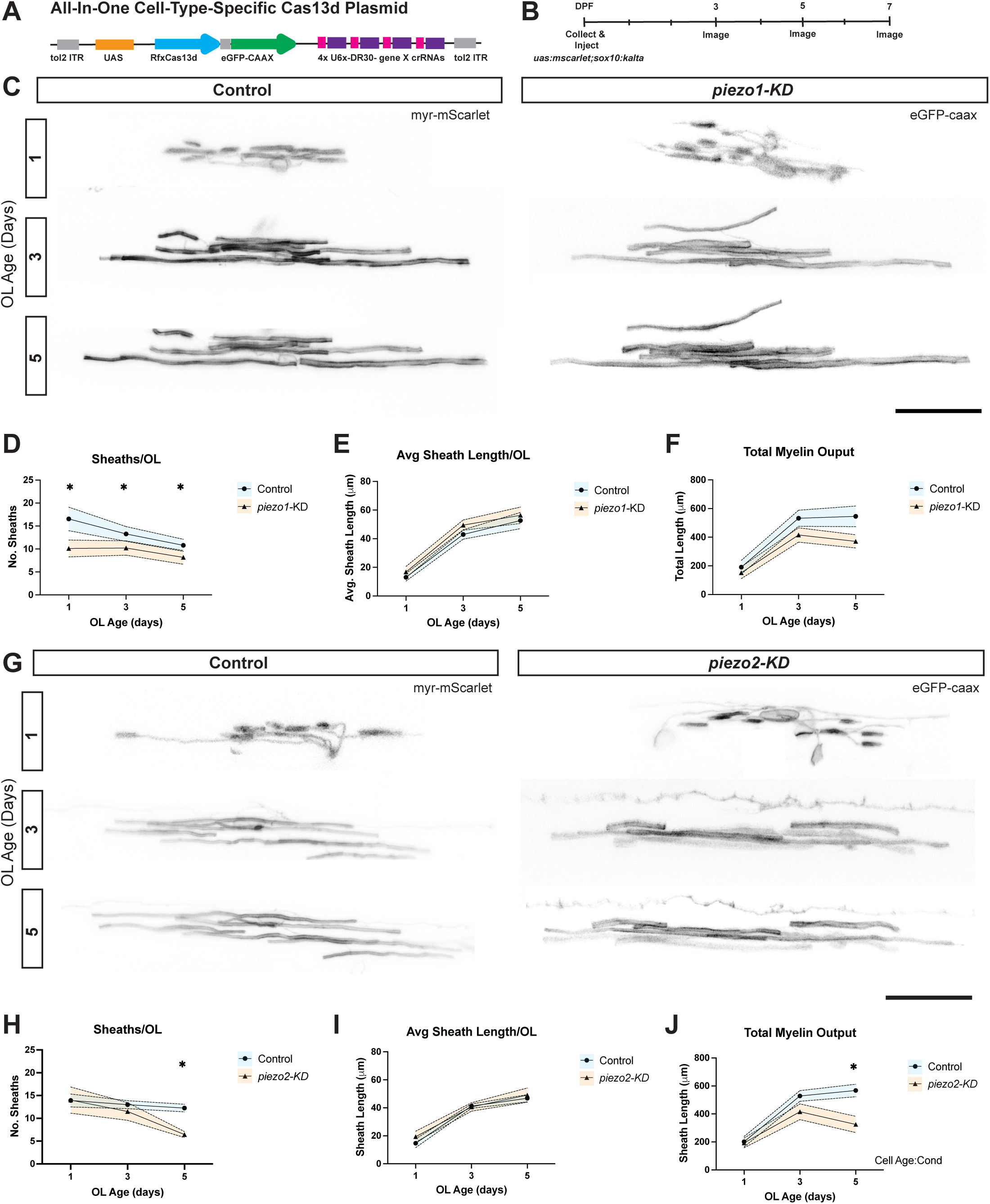
Independent targeting of *piezo1* and *piezo2* suggests distinct functions within the OL lineage. A. Schematic displaying the features of the knockdown Cas13d targeting system. Tol2 allows integration into the genome, eGFP-CAAX enables morphological characterization of cells, and 4 sites for crRNA guides mediate gene-specific knockdown. B. Experimental timeline starting with embryo harvest on the first day of the experiment, followed by longitudinal imaging of zebrafish embryos at 3, 5, and 7 dpf. C. Representative images of non-targeted control and *piezo1-KD* cell tracked through time from OL age 1-5, with day 1 being the first day of sheath formation. D. Number of sheaths per OL for non-targeted control and *piezo1-KD* cells across the experimental timeline. Control: OL1/3: N = 13 cells; 9 fish. OL5: 7 cells; 5 fish. *piezo1-KD*: OL1/3: N = 12 cells; 7 fish OL5: 9 cells; 6 fish. LMEM (Linear Mixed Effects Model) p_cond_: 0.002 p_cell age:cond_:0.009 t-test p_OL1_: 0.024, p_OL3_: 0.059, p_OL5_: 0.039 E. Average sheath length per OL for non-targeted control and *piezo1-KD* cells. Control: OL1/3: N = 13 cells; 9 fish. OL5: 7 cells; 5 fish. *piezo1-KD*: OL1/3: N = 12 cells; 7 fish OL5: 9 cells; 6 fish. LMEM p_cond_: 0.121 p_cell age:cond_: 0.248 t-test p_OL1_: 0.448, p_OL3_: 0.194, p_OL5_: 0.665 F. Total myelin output per OL for non-targeted control and *piezo1-KD* cells. Control: OL1/3: N = 13 cells; 9 fish. OL5: 7 cells; 5 fish. *piezo1-KD*: OL1/3: N = 12 cells; 7 fish OL5: 9 cells; 6 fish. LMEM p_cond_: 0.123 p_cell age:cond_:0.234 t-test p_OL1_: 0.545, p_OL3_: 0.135, p_OL5_: 0.052 G. Representative images of non-targeted control and *piezo2-KD* cells tracked from OL age 1-5, with day 1 being the first day of sheath formation. H. Number of sheaths per OL for non-targeted control and *piezo2-KD* cells. *Control*: OL1/3: N = 13 cells; 9 fish. OL5: 12 cells; 8 fish. *piezo2-KD*: OL1/3: N = 14 cells; 10 fish OL5: 9 cells; 8 fish. LMEM p_cond_: 0.046 p_cell age:cond_: 0.371 t-test p_OL1_: 0.981, p_OL3_: 0.456, p_OL5_: <0.001 I. Average sheath length per OL for non-targeted control and *piezo2-KD* cells. *Control*: OL1/3: N = 13 cells; 9 fish. OL5: 12 cells; 8 fish. *piezo2-KD*: OL1/3: N = 14 cells; 10 fish OL5: 9 cells; 8 fish. LMEM p_cond_: 0.207 p_cell age:cond_: 0.377 t-test p_OL1_: 0.364, p_OL3_: 0.850, p_OL5_: 0.669 J. Total myelin output per OL for non-targeted controls and *piezo2-KD* cells. *Control*: OL1/3: N = 13 cells; 9 fish. OL5: 12 cells; 8 fish. *piezo2-KD*: OL1/3: N = 14 cells; 10 fish OL5: 9 cells; 8 fish. LMEM p_cond_: 0.634 p_cell age:cond_: 0.033 t-test p_OL1_: 0.864, p_OL3_: 0.115, p_OL5_: 0.003 For C/G – Scale bar: 25 μm. For E-F and H-J graphs are mean ± SEM and * represent statistically significant timepoints.

### Knockdown of *piezo2* results in a mild reduction in the number of sheaths per OL and total myelin output that emerges over time

Although our RT-qPCR from primary rat OLCs did not detect *Piezo2* expression, snNucSeq and bulk sequencing from our group and others have shown *Piezo2* expression in mature OLs isolated from the optic nerve and cortex of mice^16,17,38,18^. It is likely that the variation in reported expression is the result of differences in isolation or sequencing techniques used between studies, and that *Piezo2* is indeed expressed in OLCs, making these cells one of the few cell types to express both members of the Piezo family of ion channels^24^. PIEZO1 and PIEZO2 respond to mechanical stimuli and feature similar activation thresholds, raising the potential of compensation from *piezo2* in *piezo1-KD* OLCs in the zebrafish experiments. To begin to address this, we first wanted to determine whether independent knockdown of *piezo2* impacts myelination in the zebrafish spinal cord.

The same experimental paradigm used for *piezo1* was employed to target *piezo2* in OLCs. Two crRNA targeting *piezo2* were cloned into the KD plasmid, *ptol2-UAS-RfxCas13d-P2A-EGFP-CAAX;U6x-crRNA-piezo2* (**Fig. 3A**). The targeting vector was injected into *tg(uas:mScarlet;sox10:kalta4)* zebrafish, again allowing for targeted and non-targeted cells to be visualized within the same animal and tracked from 3 to 7 dpf (**Fig. 3B, G**). Myelin sheaths were traced using Simple Neurite Tracer in Fiji to quantify the number of sheaths, average sheath length, and total myelin output per OL. Cells with *piezo2* KD showed a statistically significant reduction in the number of sheaths per OL (**Fig. 3H**) and total myelin output (**Fig. 3J**) that emerged only at the last developmental timepoint (OL5) when compared to non-targeted controls. No change in the average sheath length per OL was observed (**Fig. 3I**) between groups at any point in development. Our data show that knockdown of *piezo2* decreases the number of sheaths and the total myelin capacity per OL over time. Next, we wanted to test if Piezo1 and Piezo2 have the capacity to compensate for one another or work synergistically to regulate myelination.

### Combined loss of Piezo1 and Piezo2 results in reduced morphological complexity of OPCs and reduced myelogenic potential of OLs

To address potential compensation and/or synergy between Piezo1 and Piezo2, a dual targeting KD plasmid *(ptol2-UAS-RfxCas13d-P2A-EGFP-CAAX;U6x-crRNApiezos)* was utilized, which encodes two crRNAs targeting *piezo1* and two crRNAs targeting *piezo2.* We injected the KD plasmid into single-cell *tg(uas:mScarlet;sox10:kalta4)* embryos, which allowed targeted and non-targeted cells to be longitudinally imaged from 3 to 7 dpf within the same animal. Morphological complexity and cell volume were quantified to determine whether Piezo1 and Piezo2 are important for OPC morphogenesis. Double knockdown (*dKD*) OPCs within the spinal cord exhibit a significant reduction in morphological complexity compared to non-targeted control cells, as determined by Sholl analysis and the Kolmogorov-Smirnov test (**Fig. 4A, B**). 3D reconstructions were used to measure volume; *dKD* cells showed a reduced volume compared with controls (**Fig. 4C, Video 3**). These data suggest that the ability of OPCs to sense and respond to mechanical force via Piezos is an important regulator of OPC volume and morphological complexity. We next wanted to examine whether double KD alters differentiation and myelination of OLCs *in vivo*.

**Figure 4:**
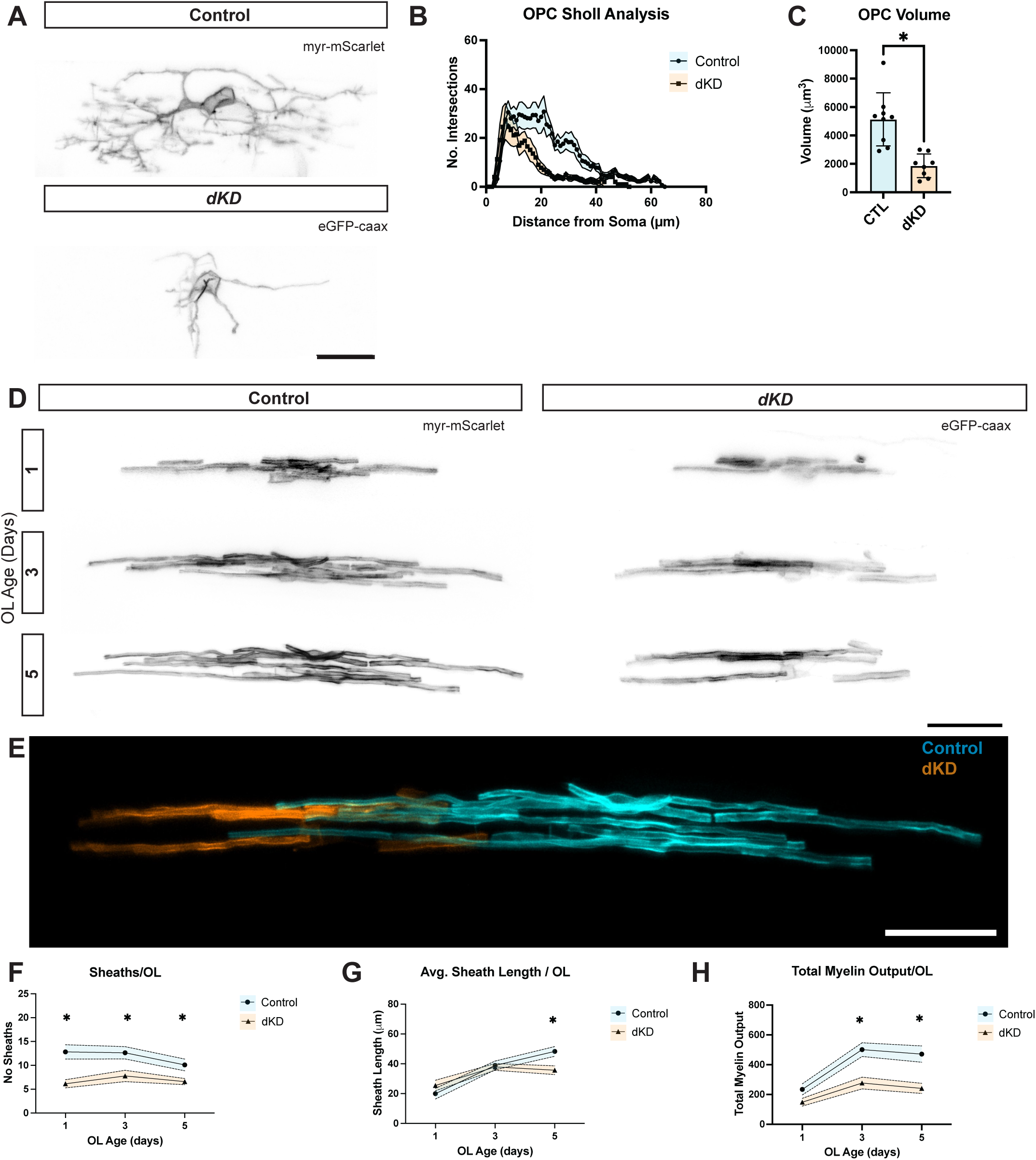
Dual targeting of Piezo1 and Piezo2 reduces OPC complexity and decreases OL sheath number, length, and total myelin output. A. Representative images of non-targeted control (myr-mScarlet) and double knockdown (dKD; eGFP-CAAX) OPCs identified in the zebrafish spinal cord. Scale Bar: 25 μm. B. Sholl analysis of reconstructed non-targeted control and dKD OPCs. Graph is mean ± SEM. Kolmogorov-Smirnov Test. Control: N = 9 cells, 7 fish; *dKD*: N = 8 cells, 6 fish; p = 0.016. C. Volume of non-targeted controls and dKD OPCs. *Control*: N = 9 cells, 7 fish; *dKD*: N = 8 cells, 6 fish; unpaired, 2-tailed, t-test p<0.001 D. Representative images of non-targeted control (myr-mScarlet) and dKD (eGFP-CAAX) cells that were tracked from OL cell age 1-5, with day 1 defined as the first day of observed sheath formation. Scale Bar: 25 μm. E. Cells tracked in Fig. 4D were developing adjacent to one another within the spinal cord and tracked from OL1 to OL5. Representative z-projection to visualize targeted and non-targeted cells within the same animal. Scale Bar: 50μm F. Quantification of the number of sheaths per OL for non-targeted control and dKD cells across the experimental timeline. Control: OL1/3: N =16 cells; 14 fish. OL5: 11 cells; 11 fish. *dKD*: OL1/3: N = 14 cells; 11 fish OL5: 10 cells; 8 fish. LMEM p_cond_: <0.001 p_cell age:cond_: 0.085. t-test p_OL1_: 0.001, p_OL3_: 0.012, p_OL5_: 0.025 G. Quantification of the average sheath length per OL for non-targeted control and dKD cells. Control: OL1/3: N =16 cells; 14 fish. OL5: 11 cells; 11 fish. *dKD*: OL1/3: N = 14 cells; 11 fish OL5: 10 cells; 8 fish. LMEM p_cond_: 0.114 p_cell age:cond_: 0.015. t-test p_OL1_: 0.298, p_OL3_: 0.780, p_OL5_: 0.010 H. Quantification of the total myelin output per OL for non-targeted control and dKD cells across the experimental timeline. Control: OL1/3: N = 16 cells; 14 fish. OL5: 11 cells; 11 fish. *dKD*: OL1/3: N = 14 cells; 11 fish OL5: 10 cells; 8 fish. LMEM p_cond_: 0.189 p_cell age:cond_: 0.004. t-test p_OL1_: 0.080, p_OL3_: 0.003, p_OL5_: 0.003 For B, F-H the graphs are mean ± SEM, and * are timepoints that are statistically significant.

Using the same genetic approach, targeted and non-targeted myelinating OLs were tracked from 3 to 7 dpf (**Fig. 4D**). The power of our cell-type-specific Cas13d system is highlighted in **Fig. 4E**, showing that targeted (dKD) and non-targeted cells (wildtype) can be visualized and tracked throughout development within the same animal (**Video 4**). dKD cells showed a significant reduction in the number of sheaths per OL, average sheath length, and total myelin output across timepoints when compared to non-targeting controls (**Fig. 4F-H**). Previously, we did not observe a reduction in average sheath length per OL with independent KD of *piezo1* or *piezo2* (**Fig. 3E, I**). dKD of *piezo1/2* revealed an enhanced phenotype across all metrics compared to targeting either MSIC alone. Altogether, our data suggest that the ability to sense mechanical cues via Piezo1/2 is important for the regulation of sheath formation, expansion, and total myelin capacity of OLs.

### Loss of PIEZOs leads to reduced sheath retractions and aberrant sheath formation outside the critical window of sheath formation

During normal OL lineage development, newly formed OLs first overproduce sheaths. Individual sheaths then undergo retraction and are removed or undergo growth and refinement^39,40^. This process is extremely well characterized in the zebrafish: once an OPC begins to differentiate, it will reach a maximum number of sheaths within 5-7 hours^40^. Over the next 1-2 days, newly formed OLs go through a selection phase, which is characterized by sheath retractions. The sheaths remaining after this critical window undergo continued growth and expansion. Given that Piezos regulate OL myelin potential and sheath expansion (**Fig. 2-4**), we wanted to examine whether Piezos could also regulate sheath retractions. We used tracings of OLs from independent and double KD of *piezo1* and *piezo2* to quantify the change in the number of sheaths per cell from OL3 to OL5, which corresponds to the developmental window when the majority of sheath retractions occur^40^. Only cells that had sheaths on the first day of imaging were included to normalize for the age of the cell. Non-targeted control cells showed the characteristic overproduction of sheaths followed by a period of selection and refinement (**Fig. 5A,B**). In contrast, dKD cells showed no change in the number of sheaths per OL during the normal retraction window and occasionally displayed sheath production during this time, which was never seen in control cells. Independent KD of *piezo1* and *piezo2* had no impact on normal sheath dynamics and retractions (**Fig. 5A,B**). The change in sheath retractions that occurs between 5 to 7 dpf suggests that Piezo1 and Piezo2 work together to regulate retractions during critical window of sheath formation.

**Figure 5:**
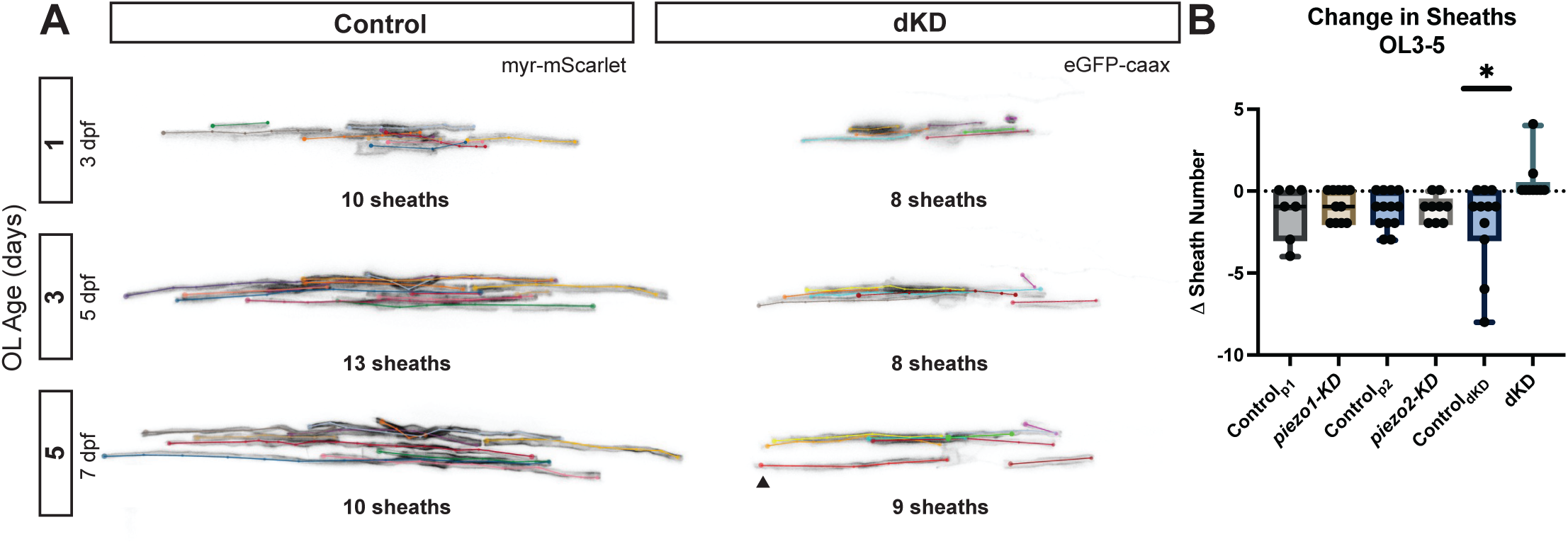
Targeting both Piezo1 and Piezo2 reduces sheath retractions and leads to aberrant sheath formation outside of the normal developmental window. A. Example tracing of OLs used to quantify the number of sheaths per OL for non-targeted control and double knockdown (dKD) cells tracked from OL1-5 (3 to 7 dpf). Arrowhead indicates the formation of a new sheath generated outside the normal developmental window. B. Change in the number of sheaths per OL between 5 to 7 dpf for each condition. The targeting of *piezo1*, *piezo2*, and *dKD* were performed as independent experiments; therefore, corresponding control groups are shown for each experiment. *Control_P1_*: N = 7 cells, 7 animals; *piezo1-KD*: N = 11 cells; 8 fish; p = 0.536 *Control_P2_*: N = 13 cells, 10 animals; *piezo2-KD*: N = 9 cells, 8 fish; p = 0.781 *Control_dKD_*:11 cells, 11 fish; *dKD:* 10 cells, 8 fish; p = 0.014

## DISCUSSION

OPCs and OLs are exposed to a range of environmental mechanical forces throughout their lifespan^41^. Our work expands the understanding of how OLCs sense and respond to their environment. First, using whole-cell patch clamp electrophysiology, we show that OLs can sense and respond to sub-micron changes in membrane displacement. The MA current profile resembles properties of PIEZO-mediated currents in other cell types. In combination with expression and protein abundance, we hypothesized that PIEZO1, a nonselective cationic MSIC, contributes to the mechanosensitivity of OPCs. Although loss of *Piezo1* does not alter proliferation or differentiation in primary mouse OLCs, we show that abolishing the ability to sense mechanical cues via PIEZO1 is sufficient to alter the cell-autonomous regulation of sheath length and total myelin capacity of OLs using inert nanofibers *in vitro*. *In vivo*, independent targeting of *piezo1* and *piezo2* in zebrafish each shows relatively mild phenotypes. KD of *piezo1* alone results in a reduction in the number of sheaths per OL, while targeting of *piezo2* results in a reduction in the number of sheaths per OL and total myelin output that emerges over time. Targeting both MSICs in OLs enhances these effects, suggesting the proteins may be able to partially compensate for each other. Additionally, dKD cells also displayed altered sheath dynamics and sheath formation outside the typical developmental window. Together, these observations suggest that the ability to sense mechanical cues from the environment via PIEZOS plays an important and evolutionarily conserved role in the regulation of myelination *in vitro* and *in vivo*.

MSICs are ideal candidates to serve as intermediaries between mechanical cues from the environment and intracellular calcium events, which have been shown to regulate OLC morphology^42,10,43^. Our electrophysiology data show that OPCs have the capacity to detect mechanical cues with extreme sensitivity (**Fig. 1A-C**), which may explain the capacity of OLCs to fine-tune features of myelination to targeted axons and their surroundings. This is consistent with the recent findings of others that OLCs are mechanosensitive^22,27^. OLCs may integrate signals from the microenvironment and modulate calcium signaling in response. The combination of current properties determined from electrophysiology recordings, the expression of candidate mechanosensors, and PIEZO1 protein abundance in OPCs suggests it is likely responsible for the MA currents we recorded (**Fig. 1D-F**). PIEZO1, PIEZO2, and TMEM63A appear to have distinct and independent functions within the OL lineage based on their expression profiles and the phenotypes observed in cell-type-specific knockout experiments. PIEZOs exhibit rapidly inactivating responses, allowing PIEZOs to serve as “event detectors,” sensitive to how quickly and how much the force changes. The ability of PIEZOs to act as dynamic sensors of mechanical force supports the idea of a role in adaptive remodeling, which could be important during dynamic remodeling of OPC processes, sheath formation, and myelin expansion. In contrast to PIEZO-mediated MS currents, TMEM63A-mediated MA currents persist throughout the length of the entire stimulus^44^. TMEM63A, which is expressed at later stages of OL differentiation^20^, may be well suited to encode information about alterations in the steady state, such as the growth of an underlying axon or changes in the environment that persist over time. Although *Piezo2* expression was difficult to detect in cultured cells early into differentiation, many RNAseq datasets detect expression in mature OLs^16–18^. PIEZO2 exhibits channel properties similar to those of PIEZO1^8^, making it more difficult to deconvolve their roles in the OL lineage. In summary, MSICs have evolved to respond to different types of mechanical forces, allowing cells to detect and interpret the diverse mechanical cues they encounter throughout their lifecycle. OLCs appear to shift the expression of various MSICs, providing the ability to sense and respond to a range of mechanical forces.

Cell-type-specific knockdown experiments in zebrafish allow us to identify the independent and combined contributions of Piezo1 and Piezo2 to the regulation of the OL lineage. Based upon a reduction in the number of sheaths per OL throughout development in *piezo1-KD* cells (**Fig. 3D**), Piezo1 is required to generate the proper number of sheaths per OL. KD of *piezo2* also decreased the number of sheaths per OL and total myelin output (**Fig. 3H**), but only at the latest developmental timepoint observed (OL5), suggesting Piezo2 may regulate later stages of sheath formation or sheath stability over time. Independent KD of *piezo1* and *piezo2* suggests that each gene has independent functions within the OL lineage, which is supported by expression data and the difference in the timing of phenotypic presentation. However, dKD experiments showed an enhanced phenotype, regarding the number of sheaths, average sheath length, and total myelin output per OL (**Fig. 4F-H**), suggesting Piezo1 and Piezo2 may compensate for one another, and both contribute to the regulation of these features of myelination. Within Schwann cells, the myelinating cells of the peripheral nervous system, PIEZO1 and PIEZO2 were suggested to play epistatic roles in myelin formation^45^. PIEZO2 conditional knockouts showed delayed myelination, while PIEZO1 was an inhibitor of radial and lateral growth. In contrast to our findings in OLCs, double knockout of PIEZO1/2 did not enhance phenotypes in Schwann cells^45^ suggesting divergent evolutionary roles for Piezos in Schwann cells and OL lineage cells.

We find that loss of *Piezo1* does not alter proliferation or differentiation in primary mouse OLCs (**Fig. 2 G/I-K)**. However, the role of PIEZO1 in modulating OPC proliferation and differentiation appears to be highly context dependent. Segal and colleagues previously found that increased tissue stiffening, as occurs during normal aging or in disease, suppresses OPC proliferation and differentiation in a PIEZO1-dependent manner^25^. Although we observed no difference in the ability of *Piezo^CKO^* OPCs to proliferate or differentiate relative to control cells, our *in vitro* studies were performed in two-dimensional cultures grown on PDL-coated coverslips which may not provide the relevant physical cues to drive PIEZO activity. Congruently, *Piezo^CKO^* OLs failed to extend sheaths to the same extent as control cells when cultured on nanofibers, indicating that the presence of mechanical cues is required to uncover deficits in the *Piezo1^CKO^*cells (**Fig. 2L-O**). While *in vitro* systems are valuable for addressing cell-autonomous questions, they lack many of the intricate cell-to-cell interactions and external cues present *in vivo*.

We observed distinct effects of PIEZO1 loss on sheath formation across our *in vitro* and *in vivo* models, which is likely due to inherent differences between our experimental systems. In our *in vitro* nanofiber experiments, controls and *Piezo1^CKO^* cells are kept in monocultures and plated at a density that minimizes the overlap of cells for ease of analysis. There is also no competition between cells of different genotypes. In contrast, our *in vivo* KD experiments are performed using a sparse targeting strategy, resulting in a KD cell surrounded by a population of non-targeted cells. Therefore, it is possible that the reduction in the number of sheaths per OL *in vivo* could be the result of competition between KD and non-targeted control cells within the highly myelinated white matter tracts of the zebrafish spinal cord. Additionally, *in vivo,* there are additional mechanical cues and signals that may contribute to the regulation of sheath formation that do not exist in our reduced culture system. It is well established that neuronal activity influences the probability of an axon becoming myelinated^46–48^. There is accumulating experimental evidence that neuronal activity may be associated with physical changes in the axon. Axons undergo rapid, transient changes in axonal radius^49–51^, pressure^52^, and the shortening of the axon terminal upon action potential arrival^53^. Therefore, if OL lineage cells have decreased ability to sense activity in of the surrounding population of axons, it may influence the number of sheaths generated. *In vitro,* we observed no difference in the number of sheaths formed using inert nanofibers between control and Piezo1^CKO^ cells. On the contrary, within the zebrafish spinal cord, there was a reduction in the number of sheaths per OL in *piezo1-KD* cells relative to controls. Additionally, a recent publication suggests OPCs and glioma cells may sense neuronal activity via a non-canonical PIEZO1 signaling mechanism. In the presence of neuronal activity, OPCs release cleaved neuroligin-3 (NLGN3) and CSPG4. NLGN3-CSPG4 interactions lead to a change in membrane tension, which is detected by PIEZO1 on OPCs and glioma cells, allowing for the detection of neuronal activity^27^. This raises the potential that OLs can directly respond to mechanical changes in neurons that occur in response to neuronal activity, or via a noncanonical pathway that leads to a change in OPC membrane tension. In summary, the differences in sheath formation between *in vitro* and *in vivo* paradigms are likely due to inherent differences in experimental design and the presence of additional mechanical cues *in vivo*.

Strikingly, we found that in contrast to control cells, *piezo1* and *piezo2* dKD myelinating OLs in the zebrafish spinal cord showed no sheath retractions from 5 to 7 dpf and occasionally generated new sheaths outside the normal developmental window (**Fig. 5**). To our knowledge, this phenotype has not been reported before. The overproduction and subsequent retraction of sheaths by newly formed OLs has been reported across *in vitro*^39^ and *in vivo*^35,40^ models. This process is highly conserved and well characterized in developing zebrafish. OLCs begin differentiating and myelinating the spinal cord around 3 dpf. Once an OPC initiates differentiation, it generates all of its sheaths within 5 to 7 hours^40^. Over the following 1-2 days (5-7 dpf), newly formed OLs undergo a period of sheath selection and retraction, after which the remaining sheaths are highly stable and continue to grow and expand. Therefore, it is surprising to see aberrant sheath formation outside the standard developmental window in dKD cells, and the unchanged sheath number in most cells may suggest a general dysregulation of sheath formation and retraction. Dysregulation of sheath dynamics in the normal developmental window required loss of both Piezos; independent KD of *piezo1* and *piezo2* had no impact on normal sheath dynamics and retractions. The change in sheath retractions that occurs between 5 to 7 dpf suggests that Piezo1 and Piezo2 may be working together to regulate sheath retractions and the critical window of sheath formation.

Within a living organism, cells are constantly exposed to a range of mechanical forces. Forces that activate MSICs include substrate stiffness, shear stress, tissue compression, membrane curvature, and changes in membrane tension^54,55^. Work from our group and others established that the MSICs Piezo1, Piezo2, and Tmem63a are playing an important role in shaping OL physiology. The challenge remains to identify the mechanical cues that each respective ion channel is responding to in the OLCs lifecycle given limited number of techniques available to mimic physiologically relevant mechanical forces *in vitro.* Myelination can be oversimplified into four broad steps: (1) OPC target identification, (2) initiation of contact, (3) wrapping, and (4) compaction^1^. After an OPC initiates contact with an axon, these physical connections persist throughout the myelination process and may serve as sites of information exchange. Axon diameter is positively correlated with the probability of an axon becoming myelinated and internode length^56,26^. Previous work has established that there are differences in membrane tension at OLC contact points with axons of varying diameters^57^. This difference in tension may serve as a mechanical cue that is detected by PIEZOs and influence sheath expansion, which is supported by the reduced average sheath length and total myelin output seen in *Piezo1^CKO^* cells plated onto inert nanofibers *in vitro* (**Fig. 2L-O**). This claim is further augmented by recent findings that sheath length scales with axon diameter and is detected by PIEZO1 in mice and primary cell culture^26^. Additionally, as myelination proceeds, the leading edge of the myelin sheath advances around the axon; existing layers must be displaced through mechanical force to allow for the addition of new wraps. Therefore, mechanical cues play the most prominent role in myelination during OPC target selection and during sheath expansion. This is consistent with the reduction in the sheath number, average sheath length and total myelin output observed in dKD cells *in vivo* (**Fig. 4D-H**). These data suggest that PIEZOs contribute to the regulation of sheath expansion and total myelin capacity of cells *in vitro* and *in vivo.* Future experiments should aim to determine where PIEZOs are localized in OLCs *in vivo* and dissect a causal link between PIEZO-mediated calcium events and sheath expansion.

## METHODS

### Isolation and expansion of OL lineage cells

Rat (**Fig. 1E-G**) or mouse (**Fig. 2**) OPCs were isolated via immunopanning from P6-9 pups as previously described^58^. Briefly, dissociated cells were passed over Ran2 and anti-GalC antibody-coated plates to deplete astrocytes and oligodendrocytes before the OPCs were positively selected for using the O4 monoclonal. OPCs from a single rat brain were expanded in poly-d-lysine (PDL) coated 3x 175cm^3^ flasks and mouse cells from a single mouse brain were expanded in a 1x 75cm^3^ dish. Cells were grown in DMEM-SATO medium containing 5 µg/ml insulin (Sigma I6634) 1 ng/ml NT-3 (Peprotech 450-03), 1 ng/ml CNTF (Peprotech 450-02) and 10 mg/ml PDGF-AA (Peprotech 100-13A). Media for mouse cells also included B27 (Gibco 17504044) to support cell health. Independent OPC isolations were performed for each trial of the *in vitro* experiments.

### Whole-cell patch clamp electrophysiology

Extracellular solution: NaCl 133 mM, KCl 3 mM, CaCl_2_ 2.5 mM, MgCl_2_ 1mM, HEPES 10mM, pH adjusted to 7.3 with CsOH, then osmolarity adjusted, if necessary, with Mannitol solution to 310 mOsm. Intracellular solution: CsCl 133 mM, EGTA 5 mM, CaCl_2_ 1 mM, HEPES 10mM, pH adjusted to 7.3 with CsOH, then osmolarity adjusted, if necessary, with Mannitol solution to 300 mOsm. Prior to use, Mg-ATP was added at 4 mM and Na-GTP was added at 0.4 mM. Patch pipettes were made from borosilicate glass O.D.:1.4 I.D.: 0.86 mm pulled on a Sutter P-97 puller and polished with a microforge MF-830. Whole cell patch clamp: Currents were recorded on cultures maintained in proliferation media in whole-cell voltage clamp mode using Axopatch 200b amplifier (Molecular Devices) sampled at 10 Hz and filtered at 2 kHz. Pipettes were used between 1.7-4.5 MΩ. Indentation and stimulation was done using a blunt glass probe positioned at ∼80° with a piezo-electric crystal microstage (E625 LVPZT Controller/Amplifier; Physik Instrumente). Cells were held at -80 mV, and indentation steps were applied at an increment of .4 µm for 300 ms.

### Fura2 Calcium Imaging

Fura2 (F1201, Thermo Fisher) was prepared as described by the manufacturer. *Piezo1*^CKO^ (*Piezo1^F/F^; Olig2^w/cre^)* or control (*Piezo1^wt/wt^; Olig2^w/cre^)* littermate control cells on coverslips were incubated in Fura2 for 15 minutes before imaging. During imaging, a ratio of 340/380 is generated, representing the emission spectra of Fura2 in its calcium-bound and unbound states. Cells were exposed sequentially to DMSO, 30 µM Yoda1, and 200 mM KCl with vacuum removal between solutions. The representative traces are averages of all cells within an ROI on a single coverslip. Imaging is performed using Lamda DG4 (Molecular Devices) with an Orca flash camera.

### Protein and RNA isolation from OL lineage cells

Primary cultured mouse or rat OPCs were grown to confluency before being plated into 60 cm PDL-coated tissue culture dishes at a density of 300k cells for OPCs and 500k cells for mature OLs. Cells were differentiated by changing the available growth factors; replacing PDGF-AA with T3 promotes OL differentiation. RNA was harvested from OPCs or OLs using a Qiagen RNAeasy Kit as per the manufacturer’s instructions and immediately stored at -80° C. For protein isolations, plates were washed with Dulbecco’s Phosphate Buffered Saline (DPBS) before lysis with 350 μL of RIPA buffer (40 mM Tris-HCL pH8.0, 150 mM NaCl, 1% NP-40, 0.5% sodium deoxycholate, 0.1% SDS, and 1mM EDTA, 0.5 mM EGTA) with the addition of protease inhibitors (11836153001, Roche). Lysate was spun at 13,000 x *g* at 4° C, and supernatant was transferred to a new tube and collected. Protein lysate was stored at -80° C.

### RT-qPCR

RNA was isolated as described using the RNeasy Mini Kit (74104, QIAGEN). cDNA was generated with SuperScript III First Strand Synthesis (18080400, ThermoFisher) and stored at -20° C. qPCR was performed with predesigned TaqMan qPCR probes (see below) on a QuantStudio 6 Flex Real-Time PCR System (4485691, ThermoFisher).

*Piezo1* (ThermoFisher, Rn01432593_m1)

*Piezo2* (ThermoFisher, Rn01491821_m1)

*Tmem63a* (ThermoFisher, Rn01415187_m1)

*Mag* (ThermoFisher, Rn01457782_m1)

*Cspg4* (ThermoFisher, Rn00578849_m1)

*GAPDH* (ThermoFisher, Rn0177563_g1)

### Western Blot

Lysates were boiled at 98° C in 1x Laemmli buffer for 5 min before being run on a Bis-Tris-gel (NP0335BOX, Invitrogen). Lysate was left unboiled when blotting for multi-pass transmembrane membrane proteins (PIEZO1-tdTomato). Transfer cassettes were used to transfer proteins to PVDF membranes (1PVH00010, Thermo Scientific). Transfer buffer (NP006-01, Thermo Scientific) with 10% methanol was run at 20V for 1-2h, depending on the size of the protein of interest. After the transfer was complete, blots were rinsed in 1x Tris Buffered Saline (TBS) with 0.1% Tween-20 (TBS-T) before blocking in 1x TBS-T with 5% milk powder for one hour on a rocker at room temperature. Blots were probed using antibodies targeting RFP, PDGFRA, MBP, and GFP (Table 1). All antibodies were used at 1:1000 with primary antibodies diluted in 2.5% BSA and TBS-T and incubated while gently rocking overnight at 4° C. Blots were washed in 3x in TBS-T and incubated with HRP-conjugated secondary at 1:5000 for two hours with 2.5% milk powder in TBS-T. Chemiluminescence was used for visualization on a Syngene GBox iChemiXT. As a loading control, preconjugated anti-ACTB-HRP (1:5000; A3854, Sigma) was used. Densitometric analyses were performed in ImageJ by quantifying bands relative to ACTB.

### EdU Incorporation Assay

Primary cultured mouse OPCs were isolated and grown to confluency before being plated onto coverslips in proliferation media (DMEM-SATO with NT3, CNTF and PDGF-AA) at a density of 8k cells per coverslip. Cells were allowed to recover for 24 hours before a 6.5-hour pulse with 10µM 5-ethynyl-2’-deoxyuridine (EdU). Cells were fixed with 4% PFA for 8 minutes followed by 3 washes with 1xPhosphate Buffered Saline (PBS). Coverslips were blocked for 1 hour in 0.2%TritonX in PBS with 5% Fetal Calf Serum (FCS) before overnight incubation with anti-OLIG2 (1:500; Olig2, Millipore). Cells that effectively incorporated EdU during the pulse period were detected using Click-iT Edu Cell Proliferation Kit with Alexa Fluor 647 (Thermo Fisher, C10340) as described in the product manual before being mounted with Fluoromount plus DAPI (Invitrogen, 00-4959-52). Images were acquired using a Zeiss ApoTome2 at 20x. 3 coverslips per animal were imaged, with 3 different regions of interest, and Olig2+ and EdU+ cells were counted manually within ImageJ. The experiment was analyzed using a paired t-test, with experiments being compared to age-matched littermate controls that were grown using the same reagents. Paired analysis accounts for inherent variability in growth rates between cultures.

### Differentiation Assay

Primary cultured mouse OPCs were isolated and grown to confluency before being plated onto coverslips at a density of 15k cells per coverslip in differentiation media (DMEM-SATO with NT3, CNTF and 50nM T3) and incubated for 48 hours. Cells were fixed with 4% PFA for 8 minutes followed by 3 washes in 1xPBS. Coverslips were blocked for 1 hour in 0.2% Triton-X in PBS with 5% FCS before overnight incubation with primary antibodies: chicken anti-MBP (1:500; MBP, Aves), rabbit anti-OLIG2 (1:500; Olig2, Millipore), and goat anti-PDGFRA (1:500; PDGFRA, R&D Systems). Coverslips were incubated with secondary antibodies for 2 hours before 3 washes in 1xPBS and mounting with Fluromount Plus DAPI (Thermo Fisher 00-4959-52). Cells were classified based on their immunopositivity for the different stages of development, and were analyzed using a paired t-test, with comparison to age-matched littermate control cultures. Images were acquired using a Zeiss ApoTome2 at 20x. 3 coverslips per animal were imaged, with 3 different regions of interest, and Olig2+ and EdU+ cells were counted manually using ImageJ. The experiment was analyzed using a paired t-test, with experiments being compared to age-matched littermate controls that were grown using the same reagents. Paired analysis accounts for inherent variability in growth rates between cultures.

### Nanofiber Assay

Nanofibers were washed in 70% ethanol before being PDL-coated overnight. Primary cultured mouse OPCs were isolated and grown to confluency before being plated at a density of 30k cells per nanofiber scaffold in 12-well plates in differentiation media (DMEM-SATO with NT3, CNTF and 50nM T3) and incubated with media changes every other day for 7 days before being fixed with 4% PFA for 8 min. Cells were blocked for 1 hour in 0.2% Triton-X in PBS with 5% normal goat serum before overnight incubation with primary antibodies: chicken anti-MBP (1:500; MBP, Aves) and rabbit anti-OLIG2 (1:500; Olig2, Millipore). Secondary antibodies (Thermo Fisher highly cross-absorbed Alexa Fluor conjugates at 1:1,000) were incubated for 2 hours. Nanofibers were removed from scaffolds and coverslipped on slides as previously described^34^ using Fluromount Plus DAPI (Thermo Fisher 00-4959-52).

Images were acquired at 20x on a Zeiss LSM 980 with Airyscan 2. The Simple Neurite Tracer plugin within Fiji was used to trace the sheaths of cells in *x*, *y*, and *z* dimensions. The number of sheaths, sheath length, and total myelin output were calculated. The experiment was analyzed using a paired t-test, with experiments being compared to age-matched littermate controls that were grown using the same reagents. Paired analysis accounts for inherent variability in growth rates between cultures.

### Primary Antibodies

**Table 1:** Primary antibodies used in the study Zebrafish husbandry.

| Target | Manufacturer | Cat # |
| --- | --- | --- |
| **OLIG2** | Millipore | AB9610 |
| **PDGFRA** | R&D Systems | AF1062 |
| **MBP** | Aves | MBP847979 |
| **RFP** | Rockland | 600-401-379 |

### Zebrafish husbandry

Experiments were performed at Oregon Health & Science University (OHSU) and under compliance with the institutional and ethical regulations for animal testing and research. *Tg(uas:mScarlet;sox10:kalta)* and *tg(sox10:kalta)* fish were maintained as heterozygotes. Experimental larvae were generated by outcross with *tg(sox10:kalta) x tg(uas:mScarlet;sox10:kalta)* and sorted for green heart and red eye at 2 dpf. From 5 days post-fertilization (dpf) until 21 dpf, zebrafish are on a diet of rotifers and dry food (Gemma 75). From 21 dpf to 3 months, zebrafish continue a diet of rotifers and increased size of dry food (Gemma 150). After adulthood, zebrafish are switched to a diet of brine shrimp and dry food (Gemma 300). Sexual differentiation and dimorphism have not occurred during the experimental window; therefore, sex could not be considered as a biological variable.

Zebrafish lines:

*Tg(uas:mscarlet)*^59^

*Tg(sox10:kalta)*^60^

### Mouse Husbandry

All mice were housed in OHSU animal facilities and maintained in a temperature and humidity-controlled environment on a 12h light/dark cycle. All procedures were approved by OHSU Institutional Animal Care and Use Comimittee (protocols TR02_IP01148 and TR02_IP00001328). Piezo1^F/F^ mice were purchased from Jackson Laboratories (B6.Cg-Piezo1 tm2.1Apat/J; Jax strain: 029213)^30^, and crossed to a Olig2-Cre (Olig2-TVA-IRES-Cre KIKO; Jax strain: 011103) mice^31^ which were a gift from Dr. David Rowitch. All experiments were performed comparing control *Piezo1^Wt/WT^;Olig2^Cre/+^*animals to *Piezo1^F/F^;Olig2^Cre/Wt^* experimental animals to account for any effects of having a single copy of the *Olig2* allele. Genotypes were determined via PCR using established primers for each line. The following primers were used:

P1 F: CTT GAC CTG TCC CCT TCC CCA TCA AG

P1 KO R: AGG TTG CAG GGT GGC ATG GCT CTT TTT

P1 WT/Fl R: CAG TCA CTG CTC TTA ACC ATT GAG CCA TCT C

Cre700F: AAG AAC CTG ATG GAC ATG TTC AGG GA

Cre700R: CCA GAC CAG GCC AGG TAT CTC TG

CreInt F: GTT TCC TCC TCT GAA GCA GGG TC

CreInt R: TAG CAC CAC CAG CAG CAG GTT C

### Cell-Type Specific Knockdown with Cas13d

Cas13d guides were designed and cloned as previously described. crRNA guides were designed by targeting 22-23 bp regions of RNA that are predicted to have good accessibility based upon analysis of the tertiary structure of RNA using RNAfold software. crRNAs were cloned into a targeting vector by restriction digest, before insertion via Gateway cloning into the destination vector pDest-4crRNA-UAS-Cas13d-P2A-EGFP-CAAX. For targeting of *piezo1* and *piezo2* independently, two crRNA targeting each gene and two non-targeting crRNA were used. For the double knockdown construct, 4 total crRNA were used: 2 targeting *piezo1* and 2 targeting *piezo2*.

*Tg(uas:mscarlet;sox10:kalta)* fish were outcrossed to *tg(sox10:kalta)* fish to generate embryos with sparse labeling of mScarlet+ non-targeted wildtype cells. An injection solution of 90 µg of UAS-Cas13d-P2A-EGFP-CAAX;U6x-crRNA(4) and 125 ng tol2 was microinjected into single-cell zebrafish embryos. Myelin sheaths were reconstructed using Simple Neurite Tracer in ImageJ to quantify the number of sheaths, average sheath length, and the total myelin output per OL. A Linear Mixed-Effects Model was used for analysis that accounts for genetic manipulation and cell age. Each fish and cell were assigned a unique ID to ensure that differences observed are due to our experimental manipulation and not being driven by a single animal.

LMEM: 𝑀𝑒𝑡𝑟𝑖𝑐 𝑜𝑓 𝐼𝑛𝑡𝑒𝑟𝑒𝑠𝑡 ∼ 𝑐𝑜𝑛𝑑𝑖𝑡𝑖𝑜𝑛 ∗ 𝑐𝑒𝑙𝑙*_age_* + (1 | 𝑐𝑒𝑙𝑙*_ID_*) + (1 | 𝑓𝑖𝑠ℎ*_ID_*)

crRNA_1__*piezo1* target sequence: TGACATAGGGCCCAGTGGAGAG

crRNA_2__*piezo1* target sequence: TTTGTGACCGGAGCGACTCGGAT

crRNA_1__*piezo2* target sequence: GTCTTTCCTGTTGGTGTGTATCC

crRNA_2__*piezo2* target sequence: AGTTGAAGAGGAGGTGGAAGTA

For longitudinal imaging, zebrafish were anesthetized in tricane (.16 mg/mL), mounted laterally in 1% low-melt agarose (Sigma A9414-10G) and submerged in embryo media and tricane (.16mg/mL) on a 100mm petri dish, and imaged at 3, 5, and 7 dpf. Between imaging sessions, zebrafish were unmounted from agarose, returned to a 24-well plate, and individually housed in embryo media. Zebrafish experiments were imaged on a Zeiss LSM 980 with Airyscan 2 using at 20xW dipping objective.

### Quantification and Statistical Analysis

Myelin sheaths on nanofibers (**Fig. 2L-O**) and in the zebrafish spinal cord (**Fig. 3 C-J**, **Fig. 4 D-H**, **Fig. 5**) were traced using the Simple Neurite Tracer plugin in ImageJ, which allows for pathfinding in *x, y,* and *z* dimensions and quantification of the number of sheaths, sheath length, and total myelin output per cell. Quantification of the immunopositivity of cultured cells (**Fig. 2 E-F/H-K**) was performed manually with the assistance of the Cell Counter plugin in ImageJ. For primary cell culture experiments using *Piezo1;Olig2^Cre/Wt^*, the experiments were analyzed using a paired t-test, with comparisons to age-matched littermate controls that were grown using the same reagents (**Fig. 2 E-O**). Paired analysis accounts for inherent variability in growth rates between cultures. All Cas13d experiments were analyzed using A Linear mixed-effects model (LMEM) to assess (1) the effect of the genetic manipulation (p-value condition) and (2) determine the significance of the interaction of genetic manipulation and cell age (p-value condition: cell age). We examined the significance between control and KD cells at individual developmental timepoints using repeated t-tests. Western blots were quantified by densitometry in ImageJ – all blots were normalized to background intensity, then bands of proteins of interest were compared with loading controls. All statistical analyses were performed within Prism 10 (Graphpad), except LMEM analysis, which was performed using Matlab. Figure legends indicate the statistical tests, *P* values, and sample size. All analysis was performed blinded to genotype.

## Acknowledgements

We would like to thank current and past members of the Emery and Monk labs, particularly Austin Forbes, Suhail Akram, Emma Brennan, Tia Perry, and Adriana Reyes for their support in zebrafish and mouse husbandry. Imaging work was supported by the OHSU Advanced Light Microscopy Core (ALMC, RRID: SCR_009991). This project was made possible by financial support provided by the Herbert R. & Jeanne C. Mayer Foundation and the OHSU Foundation Lacroute Fellowship to A.M.C., The Laura Fund to S.E.M, B.E., and K.R.M, and a National Institute of Neurological Disorders and Stroke grant (R01NS138188) to B.E. and K.R.M. B.E. was supported by an endowment from the Warren family.

## Contributions

A.M.C, B.E., and K.R.M. conceived of the project. M.E.B. provided critical unpublished data that shaped the course of experimental design. A.M.C. designed, performed, and analyzed all experiments with the following exceptions: D.O. and S.E.M. performed and analyzed whole cell patch clamp electrophysiology (Fig. 1A-C) and Fura2 calcium imaging experiments (Fig. 2A-C). D.H. designed and cloned the Cas13d plasmid targeting *piezo1* and *piezo2.* C.L.C. expanded the 3D EM segmentation of OPCs from Microns Consortium. B.E, K.R.M, D.H., and C.L.C. provided feedback on experimental design and analysis. A.M.C, B.E, and K.R.M. wrote the manuscript. All authors provided feedback on the manuscript and approved the submitted version.

---

**Supplement Figure 1: PIEZO1 is localized on the surface and processes of OPCs and is undetectable in OLs *in vitro*.**

A. Representative images of mouse OPCs plated on coverslips and stained with OPC marker (PDGFRA) and RFP to visualize PIEZO1-tdTOMATO. Scale bar = 25μm
B. Representative images of mouse OLs that were plated on coverslips and fixed 48 hours into differentiation. Cells were stained with OL marker (MBP) and RFP to detect PIEZO1-tdTOMATO. Scale bar = 25μm
C. Cells with intensity two standard deviations higher than the mean background intensity of wild-type cells were classified as PIEZO1+. The majority of OPCs (88.6%) showed detectable PIEZO1 expression, in contrast, it was detectable in only 7.2% of OLs at 48 hours into differentiation.

**Supplement Figure 2: OLCs were analyzed based on the time of first sheath formation.**

Schematic describing the classification of OLCs by cell age based upon the day of first sheath formation.

**Video 1: Fura2 calcium imaging with control cells in combination with the sequential application of DMSO, Yoda1, and KCl.**

**Video 2: Fura2 calcium imaging with *Piezo1^CKO^* cells in combination with the sequential application of DMSO, Yoda1, and KCl.**

**Video 3: 3D reconstruction of an OPC within Imaris.**

**Video 4: Z-stack showing control and dKD OLs that have developed adjacent to one another within the zebrafish spinal cord.**

