## Supplementary Figures for "PIEZOs regulate oligodendrocyte sheath formation, expansion, and myelination potential"

**Supplement Figure 1: PIEZO1 is localized on the processes of OPCs and is undetectable in OLs *in vitro*.**

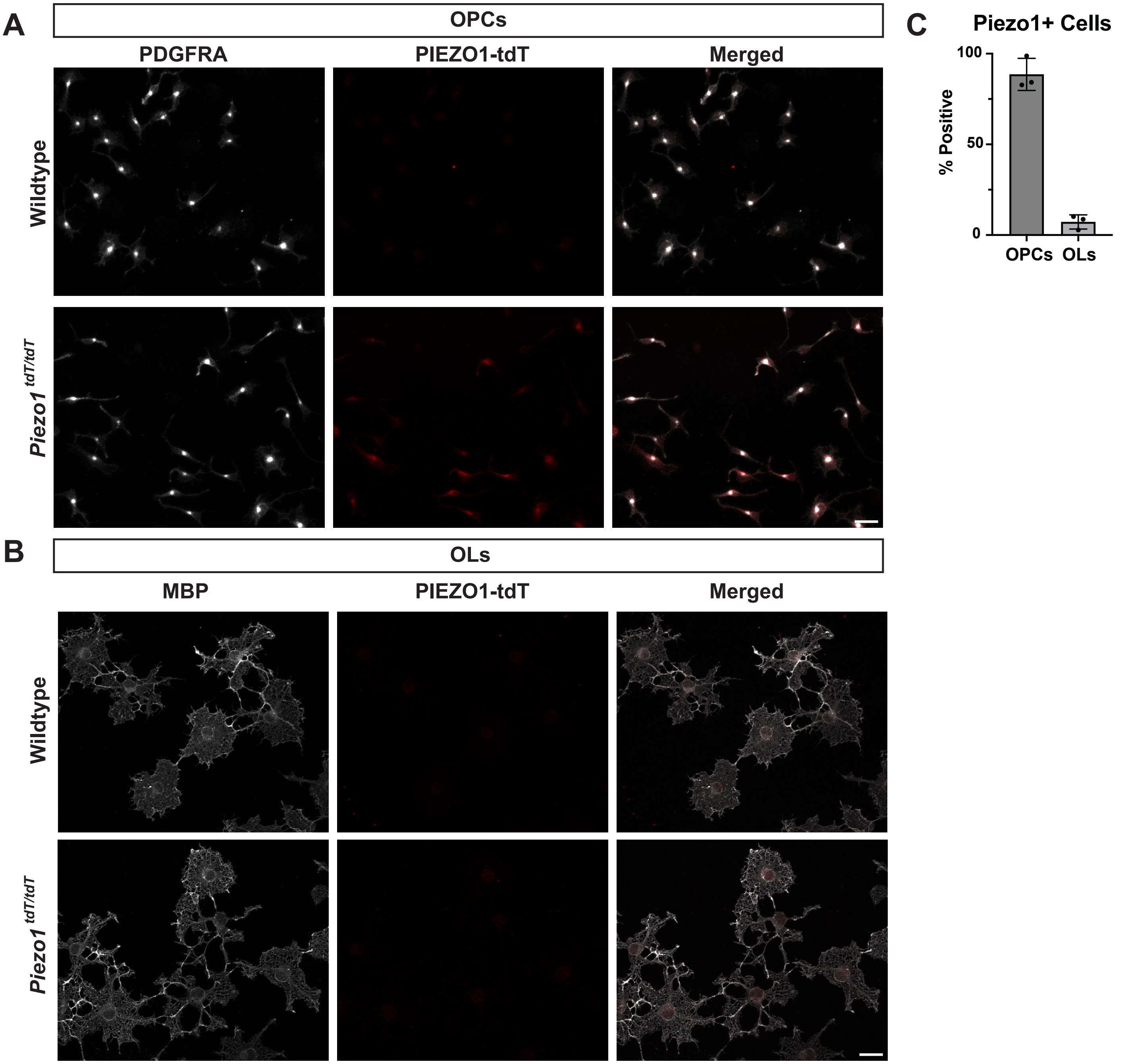

**Supplement Figure 2: OLCs were analyzed based on the time of first sheath formation.**

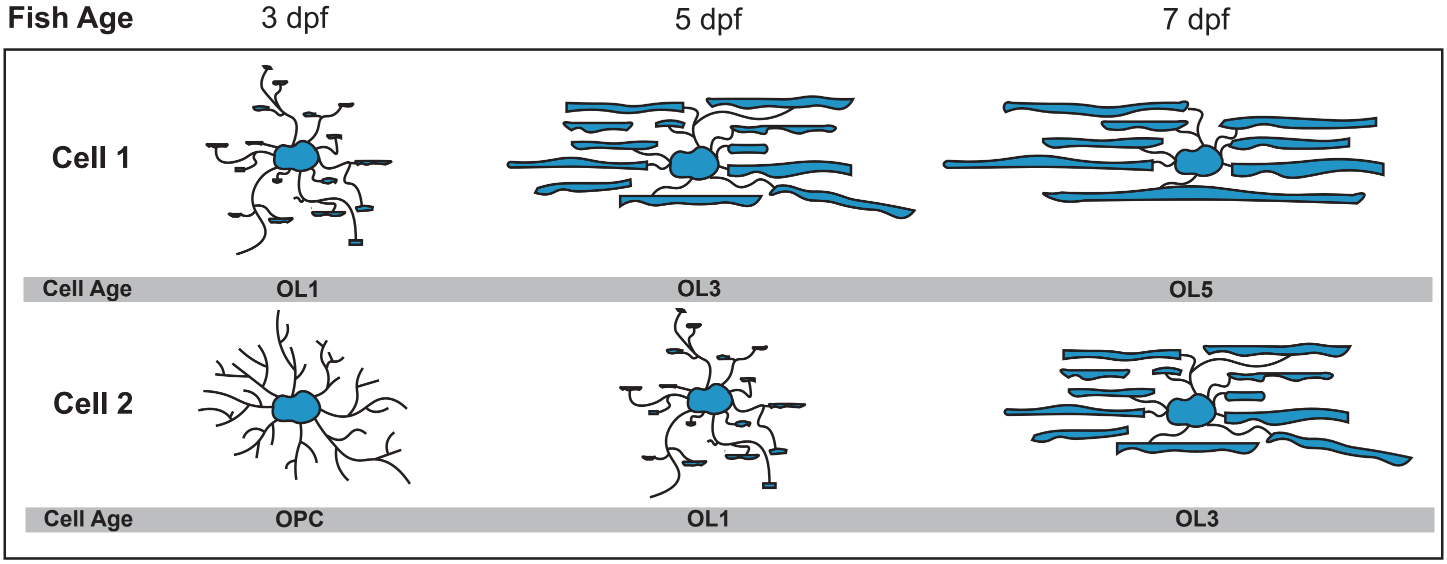
